# Design to Data for mutants of B-glucosidase B from *Paenibacillus polymyxa*: V311D, F248N, Y166H, Y166K, M221K

**DOI:** 10.1101/2023.05.10.540081

**Authors:** Morgan Chan, Justin B. Siegel, Ashley Vater

**Affiliations:** Genome Center, University of California, Davis, One Shields Avenue, Davis, California 95616, United States; Department of Biochemistry and Molecular Medicine, University of California, Davis, One Shields Avenue, Davis, California 95616, United States; Department of Chemistry, University of California, Davis, One Shields Avenue, Davis, California 95616, United States

## Abstract

Engaging computational tools for protein design is gaining traction in the enzyme engineering community. However, current design and modeling algorithms have limited functionality predictive capacities for enzymes due to limitations of the dataset in terms of size and data quality. This study aims to expand training datasets for improved algorithm development with the addition of five rationally designed single-point enzyme variants. β-glucosidase B variants were modeled in Foldit Standalone and then produced and assayed for thermal stability and kinetic parameters. Functional parameters: thermal stability (T_M_) and Michaelis-Menten constants (*k*_cat_, K_M_, and *k*_cat_/K_M_) of five variants, V311D, Y166H, M221K, F248N, and Y166K, were added into the Design2Data database. As a case study, evaluation of this small mutant set finds mutational effect trends that both corroborate and contradict findings from larger studies examining the entire dataset.

## INTRODUCTION

Enzymes can be engineered to vary reaction speed, retain activity in different reaction conditions, and catalyze novel reactions.^1-3^ Enzyme engineering has led to the development of applications including bioremediation, pharmaceuticals, and food solutions.^4,5^ However, improving in-silico functionally-targeted protein design could markedly streamline the engineering workflow. Collecting functional data on enzyme variants is an essential step to developing such algorithms.^3^ To this end, a national effort in high schools and colleges has been initiated in recent years collecting quantitative functional parameters for enzyme mutants measured in a uniform and self-consistent manner. The current model system students are working on is β-glucosidase B (BglB) from *Paenibacillus polymyxa*, a family 1 glycoside hydrolase that is structurally well characterized, biologically quite ubiquitous, and is well-suited for novice hands. Herein we describe five single point mutant BglB variants. Functional characterization data from these variants were submitted to the Design2Data (D2D) database and, in collaboration with data contributions from the D2D national network, will be used to improve enzyme design and modeling software capabilities.

The five variants selected for this study were designed in Foldit Standalone^6^, an interface of the Rosetta Molecular Modeling software package. This software evaluates protein models for structural thermodynamic stability and allows the user to examine a 3D cartoon representation of the structure that can be manipulated and modeled, engaging human spatial intuition. The five variants studied here were hypothesized to have a decrease in overall catalytic efficiency (*k*_cat_/K_M_) and thermal stability (T_M_) because Foldit predicted the mutations would reduce the enzyme’s structural thermodynamic favorability.

## METHODS

### Designing BglB mutants

Five BglB mutants were designed and modeled with Foldit Standalone^6^, a graphical user interface for the Rosetta Molecular Modeling software package. Foldit calculates a total system energy (TSE) value using the Rosetta energy score function to evaluate a protein structure.^7^ The variants included in this study had TSE score changes within 12 units of the wild type.

### Mutagenesis

Kunkel mutagenesis was carried out on BglB from *Paenibacillus polymyxa* using a standard protocol to produce mutant plasmids, which were then verified using Sanger sequencing and sequence data were analyzed in Benchling.^9,10,13^

### Protein Production and Purification

Chemically competent BLR *Escherichia coli* were transformed with mutated BglB pET 29 vectors.^7,9^ Variant expression was induced with isopropyl ß-D-1-thiogalactopyranoside (IPTG) and variants were purified through immobilized metal affinity chromatography.^9^ The total protein yield was quantified through an absorbance of 280 nm with the A280 BioTek® Epoch spectrophotometer. To confirm non-expression, variants that yielded protein concentrations of 0.3 mg/ml or less were transformed and expression and purification was replicated. Protein purity was determined with the sodium dodecyl sulfate polyacrylamide gel electrophoresis (SDS-PAGE).

### Kinetic and Thermal Stability Assays

Kinetic assays were conducted to measure the variants’ enzyme substrate conversion rates using the protocol described in previous studies.^7,9^ The catalytic efficiency of the enzyme was determined using a colorimetric assay of p-nitrophenyl-beta-D-glucoside (pNPG). Color change was observed using a spectrophotometer (A420). The kinetic constants (*k*_cat_ and K_M_) were calculated by fitting the absorbance data to the Michaelis-Menten model.^11^

The thermal stability of the protein was evaluated using a fluorescence-based Protein Thermal Shift (PTS) assay, which was carried out using the PTS kit from Thermo Fisher Scientific’s Applied Biosystem® brand.^12^ The assay was performed using a QuantaStudio 3 system at a range of temperatures from 20 to 90°C. The melting point (T_M_) was calculated by averaging the results from three technical replicates. Pearson coefficient correlation (PCC) analysis was used to evaluate the relationship between TSE and T_M_ values.^9,12^

## RESULTS

### Interpreting Foldit Models

All variants were hypothesized to exhibit a decrease in kinetic activity, decrease in thermal stability because TSE increased in the Foldit Models and all variants in this study perturbed residues in relatively close proximity to the active site. Of the five mutants, V311D was predicted to produce the smallest change in overall catalytic efficiency *k*_cat_/K_M_ due to its residues’ structural similarity (Figure 2) and its distant location from the ligand. Two mutants perturbed residue site Y166: of these two Y166H was predicted to be more thermally stable and more catalytically efficient than Y166K, due to its structural similarity to the WT residue and a smaller increase in TSE. The kinetic results supported our hypothesis in that all variants reduced *k*_cat_/K_M_, and V311D retained the most activity of the set. The thermal stability results failed to support our hypothesis, variants V311D and Y166H decreased the protein’s thermal stability, while M221K enhanced thermal stability. Y166K and F248N had reduced expression beyond the limit of expression, making it impossible for thermal stability and kinetic analysis.

**Figure 1.**
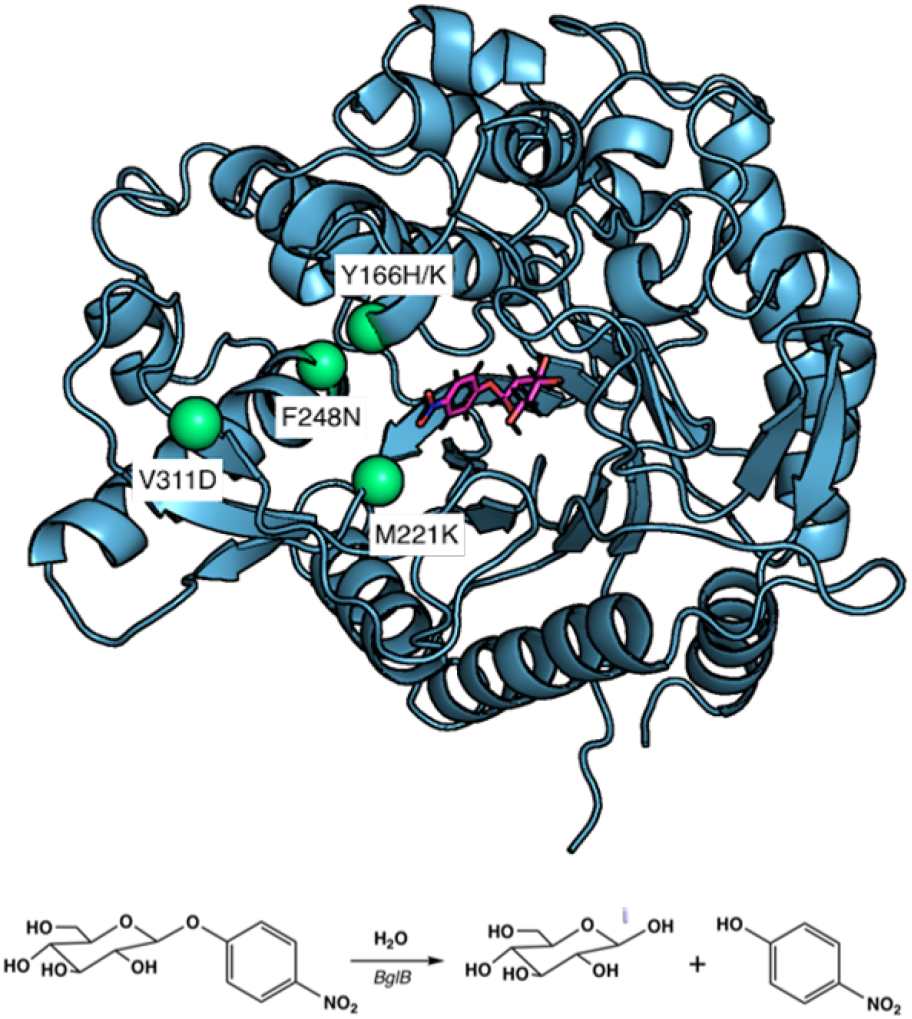
Modeled BglB-pNPG complex. PyMOL rendering^8^ of BglB with *p*-nitrophenyl-β-D-glucoside (pNPG) showing the five sequence positions selected for mutation in this study (green spheres) and the modeled transition-state structure (pink ball and stick model). Below, reaction scheme of the hydrolysis of pNPG by BglB used to determine functional T_M_ and kinetic parameters *k*_cat_, K_M_, and *k*_cat_/K_M_.

**Figure 2.**
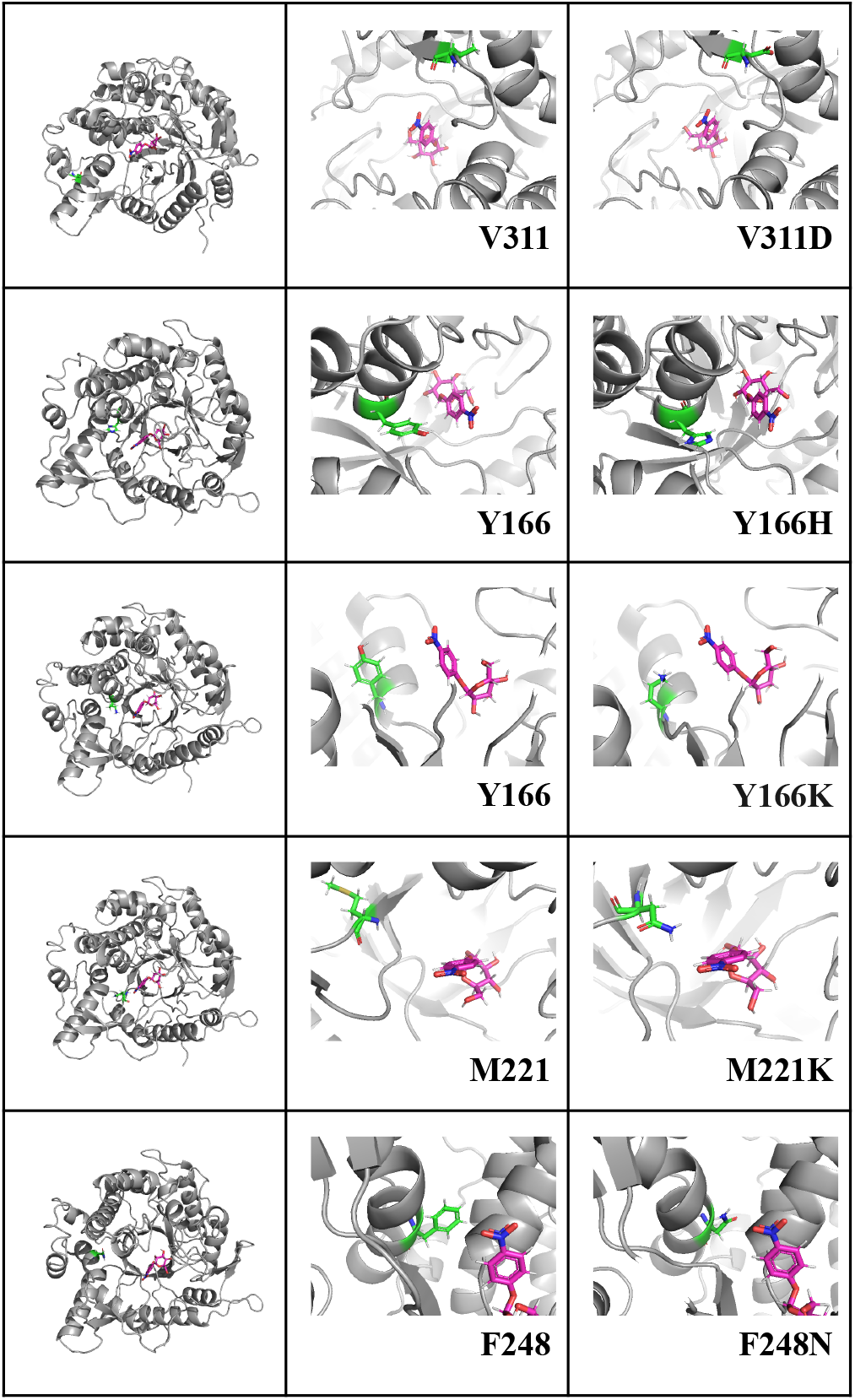
Structural analysis of Foldit models of designed BglB point mutants included in this study visualized with PyMol. Five mutant panels are shown from top to bottom by increasing TSE (except for Y166K). The point mutations (green) were shown in comparison to the ligand (pink), left-most column shows mutant location relative to the entire protein, and middle and right columns show local area of the mutations in the BglB WT protein and Foldit model, respectively.

### Protein Purity and Expression

All variants yielded protein concentration > 0.1 mg/ml, as determined by A280. SDS-PAGE protein visualization supported A280 results: Y166K, F248N, and M221K had faint to nearly invisible bands (Figure 3) and had protein yields < 0.5 mg/ml (Table 1). Mutants V311D and Y166H had more distinct bands (Figure 3) and had protein yields > 1 mg/ml (Table 1).

**Table 1.**
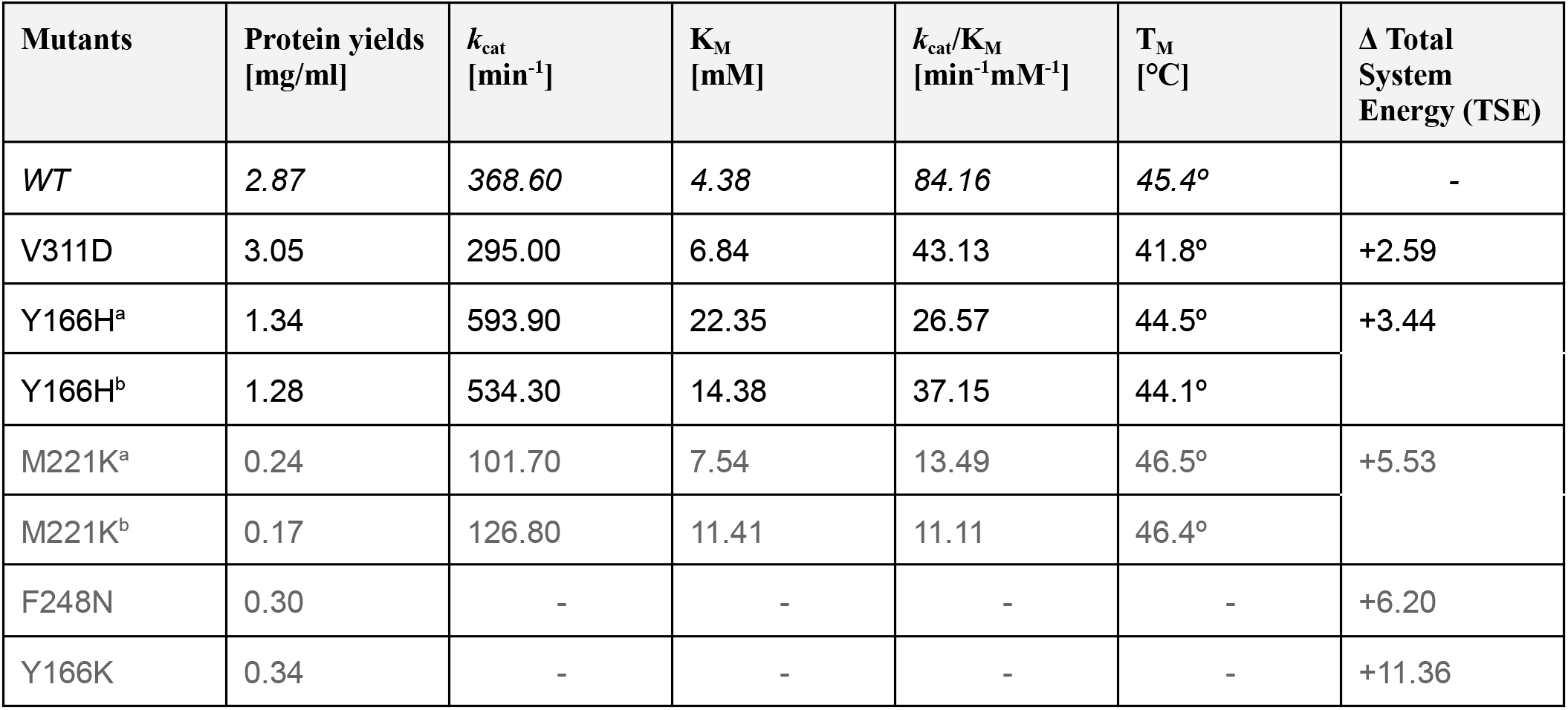
Summary of the functional parameters and model results collected from V311D, Y166H, M221K, F248N, and Y166K and BglB WT protein. *Data included biological replicates for variants Y166H and M221K. Rows are sorted by increasing change in TSE. Variants with protein yield < 0.5 mg/ml are reported in dark gray text.

| Mutants | Protein yields [mg/ml] | $k_{cat}$ [min<sup>-1</sup>] | $K_M$ [mM] | $k_{cat}/K_M$ [min<sup>-1</sup>mM<sup>-1</sup>] | $T_M$ [°C] | $\Delta$ Total System Energy (TSE) |
| --- | --- | --- | --- | --- | --- | --- |
| WT | 2.87 | 368.60 | 4.38 | 84.16 | 45.4° | - |
| V311D | 3.05 | 295.00 | 6.84 | 43.13 | 41.8° | +2.59 |
| Y166H<sup>a</sup> | 1.34 | 593.90 | 22.35 | 26.57 | 44.5° | +3.44 |
| Y166H<sup>b</sup> | 1.28 | 534.30 | 14.38 | 37.15 | 44.1° |  |
| M221K<sup>a</sup> | 0.24 | 101.70 | 7.54 | 13.49 | 46.5° | +5.53 |
| M221K<sup>b</sup> | 0.17 | 126.80 | 11.41 | 11.11 | 46.4° |  |
| F248N | 0.30 | - | - | - | - | +6.20 |
| Y166K | 0.34 | - | - | - | - | +11.36 |

| Mutants | Protein yields<br>[mg/ml] | $k_{\text{cat}}$<br>[min<sup>-1</sup>] | $K_M$<br>[mM] | $k_{\text{cat}}/K_M$<br>[min<sup>-1</sup>mM<sup>-1</sup>] | $T_M$<br>[°C] | $\Delta$ Total System<br>Energy (TSE) |
| --- | --- | --- | --- | --- | --- | --- |
| WT | 2.87 | 368.60 | 4.38 | 84.16 | 45.4° | - |
| V311D | 3.05 | 295.00 | 6.84 | 43.13 | 41.8° | +2.59 |
| Y166H<sup>a</sup> | 1.34 | 593.90 | 22.35 | 26.57 | 44.5° | +3.44 |
| Y166H<sup>b</sup> | 1.28 | 534.30 | 14.38 | 37.15 | 44.1° |  |
| M221K<sup>a</sup> | 0.24 | 101.70 | 7.54 | 13.49 | 46.5° | +5.53 |
| M221K<sup>b</sup> | 0.17 | 126.80 | 11.41 | 11.11 | 46.4° |  |
| F248N | 0.30 | - | - | - | - | +6.20 |
| Y166K | 0.34 | - | - | - | - | +11.36 |

**Figure 3.**
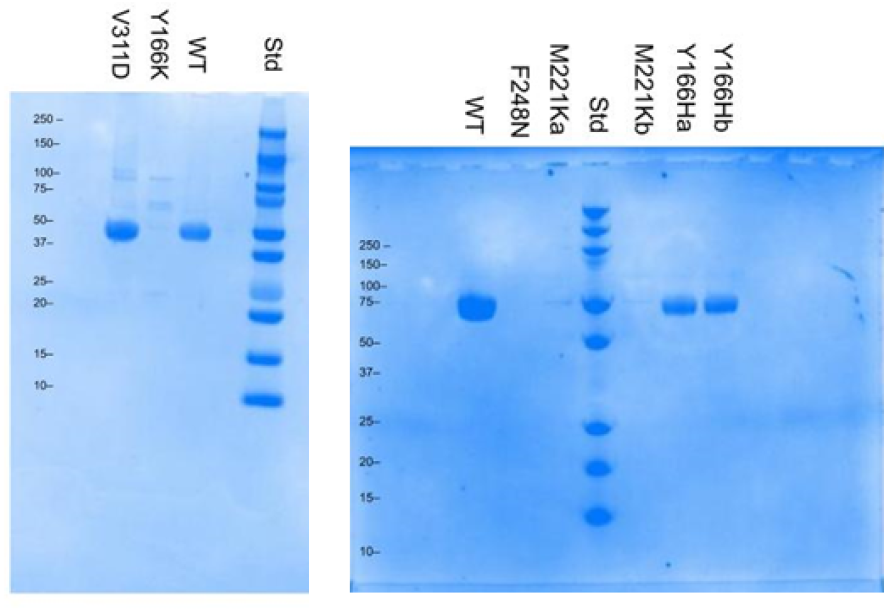
SDS-PAGE gel of mutants and two wild type enzymes. Sample bands formed at 50 kD show BglB purity. Mutants from left to right: V311D, Y166K, F248N, M221K, Y166H. Mutants F248N and Y166K have expressions below the limit of detection. M221K shows low expression, while V311D and Y166H have robustly expressed protein.

**Figure 4.**
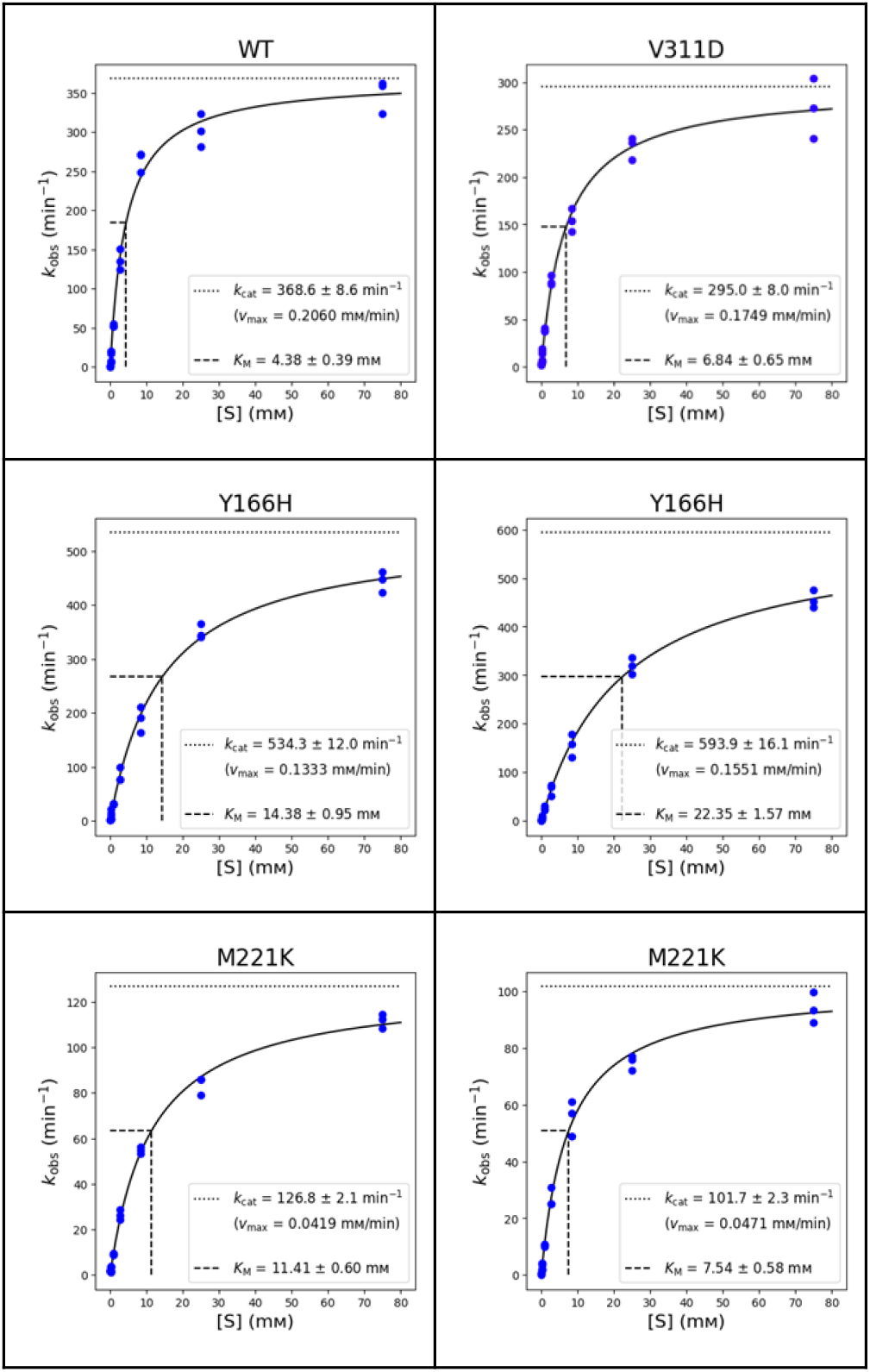
Michaelis-Menten Curves. All mutants with detectable expression and WT enzymes were assayed as technical triplicates with all data points shown in plots. Concentrations of pNPG are plotted on the horizontal axis with units of mM and kobs on the vertical axis of min^-1^. The data were fitted to the Michaelis-Menten equation through non-linear regression to determine the kinetic parameters *k*_cat_ and K_M_. The line of best-fit for the triplicate data is illustrated in each panel.

### Kinetic Activity

Colorimetric (A420) assay measured the conversion of substrate into product by monitoring the hydrolysis of p-nitrophenyl-β-D-glucoside and observing accumulation of released deprotonated nitrophenol. The WT *k*_cat_/K_M_ value was 84.16 mM^-1^ min^-1^ (Table 1), which is higher than the other mutant values. The mutant with the highest catalytic efficiency was V311D, with a 43.13 mM^-1^ min^-1^ (Table 1). Y166H had a considerable increase in the *k*_cat_ value with an average of 564.1 min^-1^, which is approximately 200 min^-1^ higher than the WT. However, the K_M_ increased approximately three-fold, negating the effects to the enhanced turnover rate. V311D showed negligible changes in their *k*_cat_ values with a value of 295 min^-1^. M221K showed markedly different changes with a value of ∼114.25 min^-1^ (Table 1). F248N and Y166K have too low expression for *k*_cat_ analysis. V311D, Y166H, and M221K had an increased K_M_ compared to the WT.

### Thermal Stability

Thermal stability was measured by thermal shift assay and melt curve data were analyzed in QuantStudio. T_M_ values were calculated by the software for each sample and averaged across technical triplicates. BglB WT protein T_M_ was 45.4 ºC (Table 1). V311D and Y166H had a lower T_M_ value, with 41.8 ºC and 44.3 ºC, respectively, while M221K had a higher T_M_ with 46.45 ºC. Figure 5 visualizes a comparison between T_M_ values for the variants and WT.

**Figure 5.**
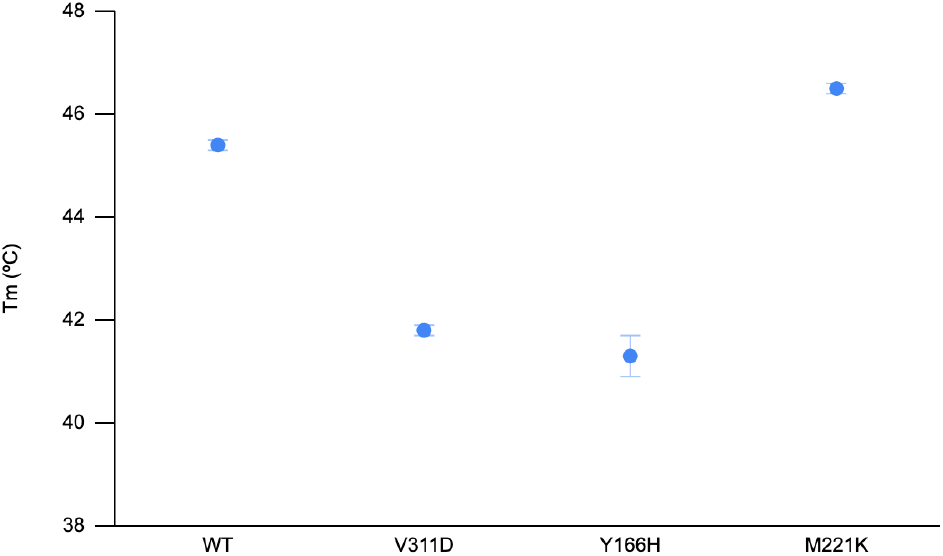
Melting temperatures of BglB mutants and wild type plotted to visualize relative thermal stability. WT, V311D, Y166H, and M221K were plotted with average T_M_ values in units of degrees Celsius. M221K and Y166H replicates were averaged. Bars indicate standard error from triplicate samples.

### Comparing Protein Solubility and Foldit Predictions

When modeled in Foldit, all 5 variants produced an increased TSE, indicating the mutations were structurally somewhat destabilizing. We evaluated the relationship between TSE and soluble expression (Figure 6) and observed a statistical significance between TSE and protein yield (p-value = 0.1141).

**Figure 6.**
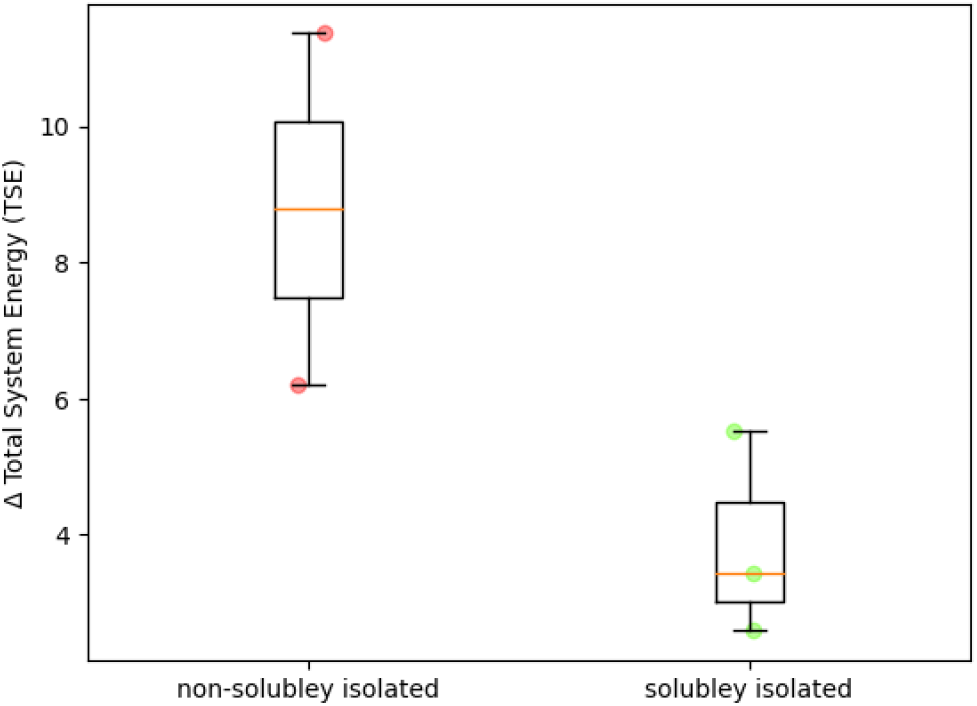
Relationship between Foldit score changes between variants and BglB WT (Total System Energy) and change in soluble isolation. Changes in TSE values compared to the WT are plotted in the vertical axis. The horizontal axis represents the differences between two groups: non-solubley isolated (Y166K and F248N) and solubley isolated (V311D, Y166H, and M221K).

## DISCUSSION

This study aimed to evaluate the effects of five single point mutations, which were chosen based on Foldit structural analysis and had a score change range of 12 TSE units. Our original hypothesis suggested that these mutations would reduce catalytic activity, and the data that we gathered supports this prediction.

Foldit TSE is accepted as a coarse indicator for likelihood of soluble protein expression.^9^ Mutations that increase the BglB WT protein TSE score by 10 units have a high likelihood to fail to express.^12^ These data support this previously observed trend.

Two variants of Y166 were included in this study and these juxtaposed, providing insights on this seemingly structurally and catalytically important site. While Y166K reduced expression beyond the limit of detection, Y166H had only a modest impact on expression levels and T_M_, but almost doubled *k*_cat_. While, any meaningful change in overall catalytic activity was negated by the 3-fold increase in K_M_, independently, the impact on each kinetic parameter provides valuable data on the structure-function relationship wherein we uncovered a site sensitive to these perturbations. The functional data supports basic structural reasoning. The wild type tyrosine and the histidine share the core heterocyclic motif. Removing the hydrophilic hydroxyl group from the distal end of the tyrosine side chain perhaps triggers cascading interactions that impact the active site dynamics and ultimately catalysis, simultaneously enhancing turnover rate and reducing binding affinity. However, when exploring a residue that is structurally and chemically very different (e.g. lysine), a four carbon chain with a charged ε-ammonium end group, we managed to break the protein. Y166 lies just at the mouth of the active site and is likely both a gatekeeper and stabilizer for the active site, after all catalytic residue E164 is nearly a next door neighbor, separated by a single proline. This case serves as a reminder that single changes in the amino acid sequence can drastically affect enzyme activity, highlighting the fragility of structural and functional properties of enzymes.

Over the last five years, our research team has published a number of similar reports written by students and typically aggregate replicate results but in presenting the disaggregated data here, we provide an example of consistent results, which indicate success of development of the robust standard operating procedures ready for novice student researchers.

The data collected from the five variants and including replicates of two variants were recorded and integrated into the Design2Data database.^14^ This dataset is being developed with the aim to improve the accuracy of computational enzyme design tools to improve the efficiency of engineering workflows for human-centered problems. Taken together with the other data generated by the D2D project, the mutants in this study provide valuable information for understanding the relationship between the structure and function of enzymes.

## ACKNOWLEDGEMENTS

This work was supported by the University of California Davis, the National Institutes of Health (R01 GM 076324-11), the National Science Foundation (award nos. 1827246, 1805510, and 1627539), and the National Institute of Environmental Health Sciences of the National Institutes of Health (award no. P42ES004699). The content is solely the responsibility of the authors and does not necessarily represent the official views of the National Institutes of Health, National Institute of Environmental Health Sciences, National Science Foundation, or UC Davis.

